# Underestimation of carbohydrates by sugar alcohols in classical anthrone-based colorimetric techniques compromises insect metabolic and energetic studies

**DOI:** 10.1101/322123

**Authors:** Mélanie J.A. Body, Jérôme Casas, Jean-Philippe Christidès, David Giron

## Abstract

Physiologically based metabolic studies usually search for easy, sensitive, and cheap techniques to rapidly estimate biological parameters such as nutrient content. Colorimetric methods to estimate carbohydrates have been extensively used (over 120,000 references). However, sugar alcohols are underestimated under conventionally used analytical conditions, in particular if using the popular van Handel method. This may lead to misinterpretations of sugar implications in biological systems. We determined the anthrone reaction with various sugar alcohols and non-alcohols individually under standard conditions (Van Handel 1985). We then manipulated the proportion of either sugar alcohols or non-alcohols in three different sugar mixtures in order to estimate the impact on the total sugar estimation. In the case of a mixture with over 50% of sugar alcohols, total sugars are underestimated by 50% when using glucose as standard.

## 1. Introduction

The need for appropriate quantification of metabolites and energy content of insects is recently growing vigorously due to the emerging field of Insects as Food and Feed (Van Huis and Tomberlin 2017). Examples range from the need to maximize proteins contents and energy levels. Also, the need to have standardized product of constant composition is a challenge of this new market. Other fields of scientific investigations requiring nutrient and energy budgets are behavioural ecology questions, multitrophic interactions and conservation biology in the prospect of climate change. Because insects can trade one source of energy for another one only (see Casas et al. 2015), complete energy budgets are needed. A comprehensive if complex framework has been recently proposed, based on dynamic energy budgets (DEB, see Llandres et al. 2015). In this context, quantifying carbohydrates is key, as it is both a large source of energy and also because it comes and is stored in different forms.

Since its introduction for the determination of carbohydrates (Dreywood 1946), anthrone in sulfuric acid has been extensively employed as a convenient and specific reagent for colorimetric estimation of a variety of carbohydrates in different materials such as food, biological fluids, wastewater, pharmaceutical products, plant or animal extracts (Body et al. 2013, 2018a in prep; Cui 2005; Foray et al. 2012; Genest and Chapman 1962; Giron et al. 2002; Green and Wade 1952; Handelsam and Sass 1956; Le and Stuckey 2016; Louis et al. 2014; Prokhovnik and Nelson 1953; Van Handel 1985; Yemm and Willis 1954). Indeed, “anthrone” and “carbohydrates” (or “sugars”) appear in more than 120,000 articles in the web of science database. This easy, sensitive, and cheap analytical method allows to produce a green colour in a quantitative manner when sugars are heated with anthrone under acidic conditions (Bailey 1958; Van Handel 1985).

Usually, glucose is used as a unique standard for carbohydrate estimation, hence the results of these assays are often presented in terms of glucose-equivalent concentrations. However, while the different reactions of anthrone with the various carbohydrates have long been known by chemists (Cui 2005), there has been no systematic investigation of the consequences for the sugar quantification in complex mixtures which may be encountered in biological materials. Indeed, different sugars analysed under the same conditions could lead to different colour intensities. For example, Yemm and Willis (1954) showed that the colour produced by the disaccharide sucrose and its constituent monosaccharides glucose and fructose – which are the most common carbohydrates – varies widely after a 2 min heating time. Indeed, when the fructose absorbance is 100 %, the sucrose absorbance is 60 % and only 20 % for glucose. This difference between the intensity of colour produced decreases when the heating time increases and this difference is nearly zero after 15 min (Yemm and Willis 1954). That is why numerous analytical methods such as the Van Handel’s technique (1985) developed to determine sugar content in mosquitoes with anthrone have a heating time of about 15-20 min to minimize the difference of absorbance between different sugars. Over this limit, the colour quickly disappears and the absorption drastically decreases. This requires scientists to choose the better trade-off between inter-sugar differential reaction and loss of coloration.

Although sugar alcohols such as glycerol, sorbitol and inositol are prevalent in many biological systems (see below), little attention has been paid to the reaction of this class of compounds with the anthrone reagent (Graham 1963). Contrary to other sugars, sugar alcohols require specific conditions to be analysed. Appropriate conditions of acid concentrations, heating time, heating temperature and wavelength detection are thus necessary (Graham 1963). These optimal conditions for sugar alcohols are 60 min heating time, at 99 °C, with 0.15 % anthrone in 100 % sulfuric acid, and detection at 500 nm. For other sugars, 15-17 min heating time, at 90-92 °C, in 0.15 % anthrone in 70 % sulfuric acid, and detection at 630 nm are the optimal conditions needed (Graham 1963; Van Handel 1985).

Due to drastically different experimental conditions for sugar alcohols and other sugars, biological sample analysis is a real problem. Indeed, the comparison between two samples may be difficult if the sugar alcohols concentration varies between treatments. In that case, overall quantities of sugars may be measured similar while they differ between experimental conditions due to changes in sugar alcohol contents. This is more common than might, at first sight, be supposed.

As sugars act both as nutrients and signalling molecules for both plants and animals, the carbohydrate content could vary in composition according to numerous factors such as species, age, nutritional status, organ, location, seasonality or time of the day. In plants, sugars are involved in diverse functions like major transporter for photoassimilated carbon (Lemoine 2000), senescence triggering processes (Wingler and Roitsch 2008), anti-freezing properties (Smallwood and Bowles 2002), salt stress response (Pommerrenig et al. 2007), and biotic and abiotic stress response (Wingler and Roitsch 2008). Sugars also have crucial roles in insects as haemolymph component and structural component of exoskeleton chitin (Wyatt 1967), phagostimulant (Bernays and Simpson 1982; Glendinning et al. 2000; Hansen 1969; Schoonhoven et al. 2005), control of diapause (Pullin 1992; Pullin and Wolda 1993), heat stress response (Salvucci et al. 2000), or cryoprotectant (Miller and Smith 1975; Wolfe et al. 1998). Because sugar alcohols have key roles in insect phagostimulation (Bernays and Simpson 1982; Glendinning et al. 2000; Hansen 1969), diapause (Pullin and Wolda 1993), and thermal tolerance (Miller and Smith 1975; Wolfe et al. 1998), they are expected to vary according to a large range of factors, including changes occurring in plants as herbivores’ food source.

A recent study reported that carbohydrates present in wastewater measured using these colorimetric methods could be under- or over-estimated due to frequent interference with coexisting compounds found in wastewater (Le and Stuckey 2016). Sugars (8 non-alcohols), quantified in glucose-equivalent, were highly variable depending on colorimetric methods used, while the 3 sugar alcohols remained undetected with all methods. However, Le and Stuckey (2016) did not investigate the reason why all sugar alcohols failed to produce any chromogen, nor the impact of variable proportions of those sugars could affect the accuracy of these colorimetric methods.

To highlight the possible bias induced, we therefore determined the specific anthrone reaction with various sugar alcohols and non-alcohols individually under standard conditions (Van Handel 1985). We then manipulated the proportion of either sugar alcohols or non-alcohols in three different sugar mixtures in order to estimate the impact on the total sugar estimation.

## 2. Materials and methods

*Chemicals* — Sugars (xylitol, sorbitol, myo-inositol, mannitol, glycerol, trehalose, sucrose, lactose, glucose and fructose) and anthrone reagent were purchased from Sigma Aldrich (Sigma Aldrich, France). The anthrone reagent consisted of 1.0 g of anthrone (Sigma Aldrich, France) dissolved in 500 mL of concentrated sulfuric acid added to 200 mL of MilliQ water (Millipore corporation).

*Calibration curves* — Colorimetric quantification of carbohydrates with anthrone reagent determines both reducing and non-reducing sugars because of the presence of the strongly oxidizing sulfuric acid. Like the other methods, it is non-stoichiometric and therefore it is necessary to prepare a calibration curve using a series of standards of known concentrations.

Stock solution of individual sugars were prepared at a concentration of 1 mg/mL in ethanol 25 %. Three sugar mixtures were also prepared with the same total sugar concentration (1 mg/mL), but using different individual sugar proportions. These artificial sugar mixtures were used compare how different sugar composition impact the accuracy of the quantification. Compositions of the three sugar mixtures were as follow: *solution 1*, 10 % sorbitol, 30 % sucrose, 30 % glucose, 30 % fructose; *solution 2*, 55 % sorbitol, 35 % sucrose, 5 % glucose, 5 % fructose; *solution 3*, 55 % sorbitol, 15 % sucrose, 15 % glucose, 15 % fructose.

*Colorimetric assay protocol* — Calibration curves were carried out for each sugar and sugar mixture following the commonly used protocol in ecology (Foray et al. 2012; Van Handel 1985). Increasing volumes (2, 5, 7.5, 10, 15, 20, 25, 30, 40, and 50 μg/μL) of 1.0 mg/mL stock sugar solution were transferred into a borosilicate tube (16 x 100 mm; Fisher Scientific, France) and placed in a water bath at 90 °C to evaporate the solvent down to a few microliters. After adding 1 mL anthrone reagent, the tubes were placed in a water bath at 90 °C for 15 min, cooled at 0 °C for 5 min, vortexed and then read in a spectrophotometer (DU®-64 spectrophotometer; Beckman, Villepinte, France) at 630 nm.

*Concentration calculation* — Sugar concentrations are usually determined using glucose standards of known concentrations. To determine the error of estimation due to the use of a single sugar calibration curve, we estimated sugar concentrations using both generic glucose standard (glucose-equivalent concentrations), and specific sugar standards (real concentrations).

## 3. Results and Discussion

The aim of this study was to characterize the anthrone reaction with sugar alcohols and other sugars and then to determine how sugar alcohols affect the colorimetric quantification of total carbohydrates in biological samples. Our results clearly show that calibration curves differ between sugars (Figure 1A). At the same concentration, all sugars do not have the same absorbance. Indeed, reaction between anthrone reagent and sugar alcohols results in lower colour intensity than for other sugars, as well as higher limit of detection (less than 2 μg/μL for non-alcohols vs. 7.5-20 μg/μL for alcohols). This observation suggests that anthrone reaction is affected by carbohydrate structure, and might be the source of some of the problems highlighted by Le and Stuckey (2016). Indeed, it could be due to the steric hindrance of sugar alcohols. The specific conformation of sugar alcohols could disturb the reagent access to some functional groups and so decrease the colorimetric reaction with anthrone.

**Figure 1.**
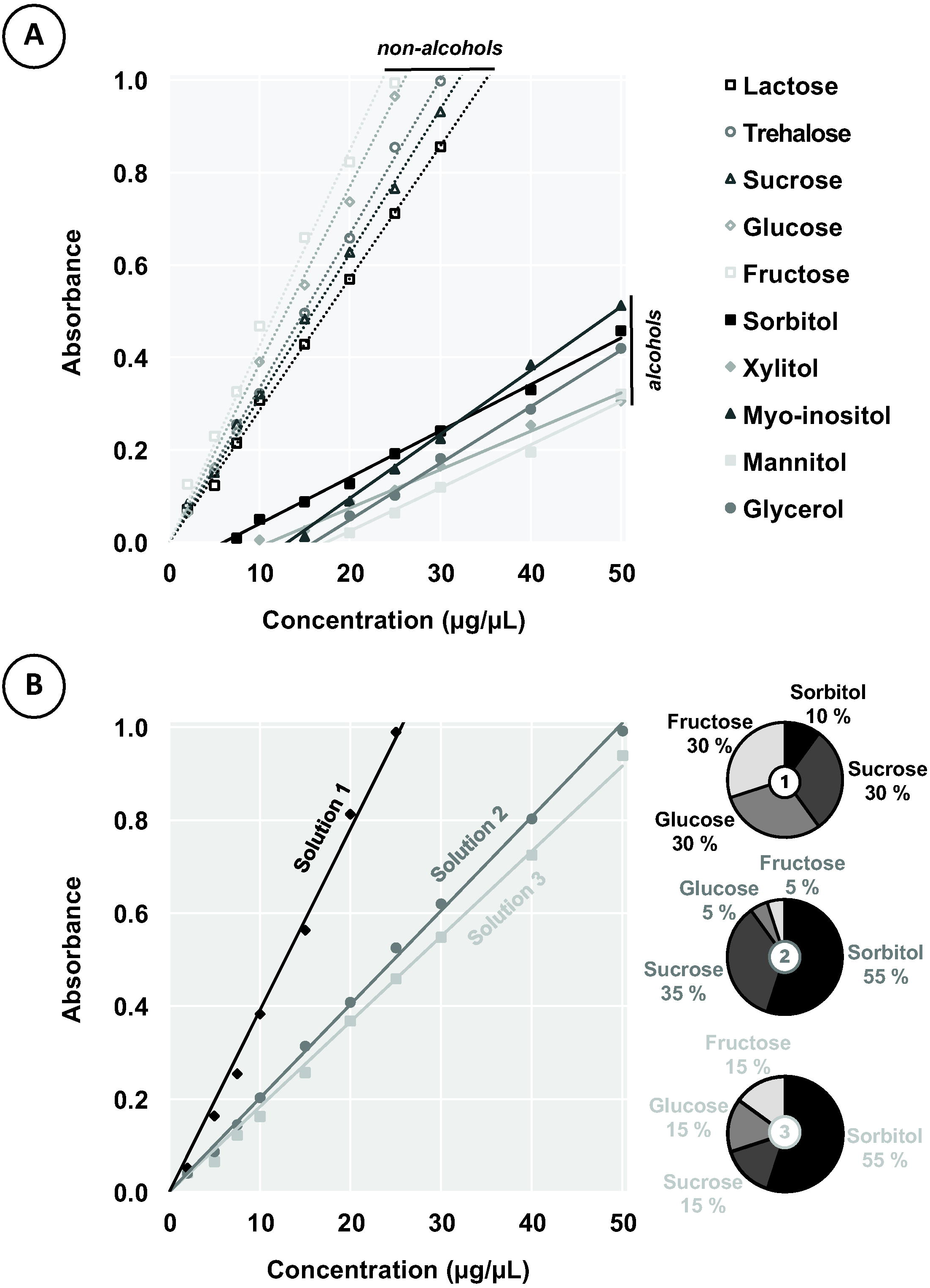
Differential reaction of sugar alcohols and non-alcohols with anthrone reagent. (A) Calibration curves for sugar non-alcohols (open markers for data points and dotted lines for linear regressions) and alcohols (closed markers for data points and solid lines for linear regressions) under Van Handel (1985) conditions which are commonly used in ecological studies. (B) Calibration curves for three sugar mixtures with the same total sugar concentration but different individual sugar proportions as presented on the chart pies. Equations and R^2^ coefficients of determination of all the calibration curves presented in this figure are summarized in Table 1.

As a consequence, we stress that the concentration of sugar alcohols has a high impact on the accuracy of the total sugar content quantification (about 80 % in the case of single sugar, Table 1). Indeed, the sugar alcohols concentration in a mixture profoundly influences the intensity of the developed colour (Figure 1B), and can lead to an underestimation of the sugar pool up to about 50 % in the case of two of our sugar mixtures (Table 1). As the sugar alcohols concentration increased, the optical density of the resulting colour decreased under standard analytical conditions (Van Handel 1985). So, in presence of sugar alcohols, standard calibration curves usually done with glucose are inappropriate and do not allow an accurate determination of the sugar content in samples. If the sugar alcohol status is modulated according to one or more factors, comparisons between samples might be biased which may impair ecological interpretations.

**Table 1.** Estimation error of sugar concentration. Equations and R^2^ coefficients of determination of all the calibration curves presented in Figure 1 are summarized here. Sugar concentrations are estimated for an absorbance of 0.5 using two different methods: the glucose standard curve (glucose-equivalent concentrations), as classically used in colorimetric assays, and a specific standard curve (real concentrations) for comparison. Specific standard curves, allowing for the calculation of real concentration, are either done using known concentration of the specific sugar to quantify (for example, using sorbitol standard to quantify an unknown concentration of sorbitol), or a mixture of sugars including the main sugars present in the solution to quantify, in similar proportions. The difference between both estimation is calculated for each sugar non-alcohol, alcohol, and mixture.

| Sugar | Equation | R<sup>2</sup> | Glucose-equivalent concentration (µg/µL) | Real concentration (µg/µL) | Difference |
| --- | --- | --- | --- | --- | --- |
| **Non-alcohol** |  |  |  |  |  |
| Trehalose | $y = 0.0334x$ | 0.9989 | 12.99 | 14.97 | -13 % |
| Sucrose | $y = 0.0312x$ | 0.9978 | 12.99 | 16.03 | -19 % |
| Lactose | $y = 0.0286x$ | 0.9981 | 12.99 | 17.48 | -26 % |
| Glucose | $y = 0.0385x$ | 0.9936 | 12.99 | 12.99 | 0 % |
| Fructose | $y = 0.0424x$ | 0.9906 | 12.99 | 11.79 | +10 % |
| **Alcohol** |  |  |  |  |  |
| Xylitol | $y = 0.0083x - 0.0922$ | 0.9921 | 12.99 | 71.35 | -82 % |
| Sorbitol | $y = 0.0100x - 0.0602$ | 0.9956 | 12.99 | 56.02 | -77 % |
| Myo-inositol | $y = 0.0138x - 0.1808$ | 0.9986 | 12.99 | 49.33 | -74 % |
| Mannitol | $y = 0.0093x - 0.1622$ | 0.9923 | 12.99 | 71.20 | -82 % |
| Glycerol | $y = 0.0122x - 0.1959$ | 0.9894 | 12.99 | 57.04 | -77 % |
| **Sugar mixture** |  |  |  |  |  |
| Solution 1<br>(10 % sorbitol, 30 % sucrose, 30 % glucose, 30 % fructose) | $y = 0.0391x$ | 0.9931 | 12.99 | 12.79 | +2 % |
| Solution 2<br>(55 % sorbitol, 35 % sucrose, 5 % glucose, 5 % fructose) | $y = 0.0202x$ | 0.9985 | 12.99 | 24.75 | -48 % |
| Solution 3<br>(55 % sorbitol, 15 % sucrose, 15 % glucose, 15 % fructose) | $y = 0.0183x$ | 0.9971 | 12.99 | 27.32 | -52 % |

For example, a study comparing sugar composition of the food source for a moth larva, photosynthetically active (green) leaves and senescing (yellow) leaves on apple-tree, reported a strong alteration of sorbitol content from 50 % in green tissue to 32 % in yellow tissue (Body et al. 2013, 2018b submitted). Such comparison, using colorimetric assays to quantify total soluble sugars, (Body et al. 2018a in prep, 2018b submitted) would be impaired by a change in sugar composition involving sugar alcohols. When quantified using glucose standards (glucose-equivalent concentrations), sugar concentrations were of 60.92 ± 20.90 μg/mg DW (average ± S.D.; DW, dry weight) on green leaves, and 43.65 ± 13.26 μg/mg DW on yellow. When a sugar mixture of the main sugars (sorbitol, sucrose, glucose, and fructose), in similar proportions as in the samples, was used for the calibration curve, instead of a single (non-alcohol) sugar, concentrations were of 107.20 ± 31.70 μg/mg DW on green leaves, and 50.62 ± 16.59 μg/mg DW on yellow leaves (real concentrations). The sugar pool was thus underestimated by 43 % on green leaves, and 14 % on yellow leaves. Such a difference in the estimation of sugar concentrations in green leaves could impact the conclusion drawn from this study, as sugar concentrations dropped only 28 % during senescing process in the first estimation, compared to a drop of 52 % between green and yellow leaves using an appropriate calibration curve. Worst estimates can be expected with increasing proportion of sugar alcohols in the sample. These changes reported in leaves could impact sugar alcohol composition of herbivores feeding on these different tissues.

As conclusion, we suggest that metabolic studies must be cautious in their interpretation of physiological data collected with such colorimetric techniques. This is particularly true in biological systems or in specific conditions where sugar alcohols are expected to be involved and we described above a wide range of situations where this happens. Alternatively, one might consider alternative techniques that allow a quantification of individual sugars (Body et al. 2013, 2018b submitted; Cui 2005; Ouchemoukh et al. 2010; Rovio et al. 2007) or, at least, a concomitant analysis of subsamples with methods adapted to either sugar alcohols or non-alcohol ones (Graham 1963; Van Handel 1985).

## Conflict of interest

The authors declare that there is no conflict of interest.

## Author contribution

MB designed the overall study, carried out the experiment, and analysed the data, with help from JPC. MB, JC, and DG wrote the manuscript.

## Funding statement

This study has been supported by the Région Centre project to DG ENDOFEED 201000047141.

## References

Bailey, R.W., 1958. The reaction of pentoses with anthrone. Biochemical Journal, 68, 669–672.

Bernays, E.A. and Simpson, S.J., 1982. Control of food intake. Advances in Insect Physiology 16: 59–118. Doi: 10.1016/S0065-2806(08)60152-6

Body, M., Kaiser, W., Dubreuil, G., Casas, J. and Giron, D., 2013. Leaf-miners co-opt microorganisms to enhance their nutritional environment. Journal of Chemical Ecology 3: 969–977. Doi: 10.1007/s10886-013-0307-y

Body, M., Behmer, S.T., Pelisson, P.-F., Casas, J. and Giron, D., 2018a in prep. Field application of the geometric framework reveals a multistep strategy of nutrient regulation in a leaf-miner.

Body, M., Casas, J. and Giron, D., 2018b submitted. Manipulation of plant primary metabolism by leaf-mining larvae in the race against leaf senescence. Ecological Monographs.

Casas, J., Body, M., Gutzwiller, F., Giron, D., Lazzari, C.R., Pincebourde, S., Richard, R., and Llandres, A.L., 2015. Increasing metabolic rate despite declining body weight in an adult parasitoid wasp. Journal of Insect Physiology 79: 27–35. Doi: 10.1016/j.jinsphys.2015.05.005

Cui, S.W., 2005. Food carbohydrates: Chemistry, physical properties, and applications. Boca Raton, FL: Taylor and Francis.

Dreywood, R., 1946. Qualitative test for carbohydrate material. Industrial and Engineering Chemistry - Analytical Edition 18: 499–499.

Foray, V., Pelisson, P.-F., Bel-Venner, M.-C., Desouhant, E., Venner, S., Menu, F., Giron, D. and Rey, B., 2012. A handbook for uncovering the complete energetic budget in insects: The van Handel’s method (1985) revisited. Physiological Entomology 37: 295–302. Doi: 10.1111/j.1365-3032.2012.00831.x

Genest, C. and Chapman, D.G., 1962. Extraction and identification of sugar alcohols and other carbohydrates in dietetic foods. Journal of the Association of Official Analytical Chemists 45: 422–424.

Giron, D., Rivero, A., Mandon, N., Darrouzet, E. and Casas, J., 2002. The physiology of host feeding in parasitic wasps: Implications for survival. Functional Ecology 16: 750–757. Doi: 0.1046/j.1365-2435.2002.00679.x

Glendinning, J.I., Nelson, N.M. and Bernays, E.A., 2000. How do inositol and glucose modulate feeding in *Manduca sexta* caterpillars? Journal of Experimental Biology 203: 1299–1315.

Graham, H.D., 1963. Reaction of sugar alcohols with the anthrone reagent. Journal of Food Science 28: 440–445. Doi: 10.1111/j.1365-2621.1963.tb00224.x

Green, P. and Wade, E., 1952. Determination of true blood sugar using anthrone. Canadian Medical Association Journal 66: 175–175.

Handelsman, M.B. and Sass, M., 1956. The determination of blood sugar by the anthrone method. Journal of Laboratory and Clinical Medicine 48: 652–659.

Hansen, K., 1969. The mechanism of insect sugar reception, a biochemical investigation. In: Olfaction and taste. Rockefeller University Press New York. 3: 382–391.

Le, C. and Stuckey, D.C., 2016. Colorimetric measurement of carbohydrates in biological wastewater treatment systems: A critical evaluation. Water Research 94: 280–287. Doi: 10.1016/j.watres.2016.03.008

Lemoine, R., 2000. Sucrose transporters in plants: Update on function and structure. Biochimica et Biophysica Acta (BBA)-Biomembranes 1465: 246–262. Doi: 10.1016/S0005-2736(00)00142-5

Llandres, A.L., Marques, G.M., Maino, J.L., Kooijman, S.A.L.M., Kearney, M.R. and Casas, J., 2015. A dynamic energy budget for the whole life-cycle of holometabolous insects. Ecological Monographs 85: 353–371. Doi: 10.1890/14-0976.1

Louis, M., Grégoire, J.C. and Pélisson, P.-F., 2014. Exploiting fugitive resources: How long-lived is “fugitive”? Fallen trees are a long-lasting reward for *Ips typographus* (Coleoptera, Curculionidae, Scolytinae). Forest ecology and management 331: 129–134. Doi: 10.1016/j.foreco.2014.08.009

Miller, L.K. and Smith, J.S., 1975. Production of threitol and sorbitol by an adult insect: Association with freezing tolerance. Nature 258: 519–520. Doi: 10.1038/258519a0

Ouchemoukh, S., Schweitzer, P., Bey, M.B., Djoudad-Kadji, H. and Louaileche, H., 2010. HPLC sugar profiles of Algerian honeys. Food Chemistry 121: 561–568. Doi: 10.1016/j. foodchem.2009.12.047

Pommerrenig, B., Papini-Terzi, F.S. and Sauer, N., 2007. Differential regulation of sorbitol and sucrose loading into the phloem of *Plantago major* in response to salt stress. Plant Physiology 144: 1029–1038. Doi: 10.1104/pp.106.089151

Prokhovnik, S.J. and Nelson, J.F., 1953. Determination of blood sugar with anthrone. Australian Journal of Experimental Biology and Medical Science 31: 279–282.

Pullin, A.S., 1992. Diapause metabolism and changes in carbohydrates related to cryoprotection in *Pieris brassicae*. Journal of Insect Physiology 38: 319–327. Doi: 10.1016/0022-1910(92)90056-J

Pullin, A.S., and Wolda, H., 1993. Glycerol and glucose accumulation during diapause in a tropical beetle. Physiological Entomology 18: 75–78. Doi: 10.1111/j. 1365-3032.1993.tb00451.x

Rivero, A. and Casas, J., 1999. Incorporating physiology into parasitoid ecology: The allocation of nutritional resources. Research on Population Ecology 41: 39–45. Doi: 10.1007/PL00011981

Rovio, S., Yli-Kauhaluoma, J. and Sirén, H., 2007. Determination of neutral carbohydrates by CZE with direct UV detection. Electrophoresis 28: 3129–3135. Doi: 10.1002/elps.200600783

Salvucci, M.E., Stecher, D.S. and Henneberry, T.J., 2000. Heat shock proteins in whiteflies, an insect that accumulates sorbitol in response to heat stress. Journal of Thermal Biology 25: 363–371. Doi: 10.1016/S0306-4565(99)00108-4

Schoonhoven, L.M., Van Loon, J.J.A. and Dicke, M., 2005. Insect-plant biology. Oxford university press.

Smallwood, M. and Bowles, D.J., 2002. Plants in a cold climate. Philosophical Transactions of the Royal Society B: Biological Sciences 357: 831–847. Doi: 10.1098/rstb.2002.1073

Van Handel, E., 1985. Rapid determination of glycogen and sugars in mosquitoes. Journal of the American Mosquito Control Association 1: 299–301.

Van Huis, A. and Tomberlin, J.K., 2017. Insects as food and feed: From production to consumption. Wageningen Academic Publishers.

Wingler, A. and Roitsch, T., 2008. Metabolic regulation of leaf senescence: Interactions of sugar signalling with biotic and abiotic stress responses. Plant Biology 10: 50–62. Doi: 10.1111/j.1438-8677.2008.00086.x

Wolfe, G.R., Hendrix, D.L. and Salvucci, M.E., 1998. A thermoprotective role for sorbitol in the silverleaf whitefly, *Bemisia argentifolii*. Journal of Insect Physiology 44: 597–603. Doi: 10.1016/S0022-1910(98)00035-3

Wyatt, G.R., 1967. The biochemistry of sugars and polysaccharides in insects. Advances in Insect Physiology 4: 287–360. Doi: 10.1016/S0065-2806(08)60210-6

Yemm, E.W. and Willis, A.J., 1954. The estimation of carbohydrates in plant extracts by anthrone. Biochemical Journal 57: 508–514.

